# High Field Asymmetric Waveform Ion Mobility Spectrometry-Mass Spectrometry to Enhance Cardiac Muscle Proteome Coverage

**DOI:** 10.1101/2022.12.28.522124

**Authors:** Lizhuo Ai, Aleksandra Binek, Simion Kreimer, Matthew Ayres, Aleksandr Stotland, Jennifer E. Van Eyk

## Abstract

Heart tissue sample preparation for mass spectrometry (MS) analysis that includes pre-fractionation reduces the cellular protein dynamic range and increases the relative abundance of non-sarcomeric proteins. We previously described “IN-Sequence” (IN-Seq) where heart tissue lysate is sequentially partitioned into three subcellular fractions to increase the proteome coverage than a single direct tissue analysis by mass spectrometry. Here, we report an adaptation of the high-field asymmetric ion mobility spectrometry (FAIMS) coupled to mass spectrometry, and the establishment of a simple one step sample preparation coupled with gas-phase fractionation. FAIMS approach substantially reduces manual sample handling, significantly shortens MS instrument processing time, and produces unique protein identification and quantification approximating the commonly used IN-Seq method in for less time requirement.

## Introduction

Modern evolution in liquid chromatography (LC) mass spectrometry (MS)-based proteomics methods have steadily increased total proteome coverage. Even so, sample fractionation prior to MS has remained a fundamental approach and permitted more thorough detection of the proteome by reducing the sample complexity [1]. Specialized proteomic sample preparation protocols [2-4] and instrumental techniques [5-7] have been developed to permit a deeper investigation into the proteome of the heart. Various studies, using drosophila, or mice, or rabbits, or human patients, have identified thousands of proteins in the cardiac proteomes, for example: [8-11].

In the heart, the sarcomeric proteins which comprise the contractile apparatus dominate the cardiac proteome [11, 12]. To increase quantification of low-abundance proteins in the cardiac tissues, sample pre-fractionation methods have been developed that take advantage of different chemical properties (e.g., In Sequence) or isolation of specific organelles or specific protein functional class. A few examples of the latter include Gramolini *et. al*., pre-fractionated mouse heart samples to “contractile” and 6 other fractions (“nuclei I”, “nuclei II”, “mitochondria I”, “mitochondrial II”, “microsome” and “cytosol fractions”) by differential ultracentrifugation in sucrose gradients before LC-MS proteomic measurement [7]. Jones *et. al*. and Boivin and Allen prepared mouse heart tissue proteins into 5 different sub-fractionations, which included additional fractions for “total membrane”, and “sarcolemmal” proteins [13-15]. Warran *et. al*., performed a series of differential centrifugations that produce organelles like “nuclear”, “mitochondrial”, “cytoplasmic”, “microsomal”, and “sarcomeric” enriched fractions during heart tissue preparations prior to LC-MS cardiac proteome analysis [16].

In the early 2000s, we used chemical properties of protein to produce a fast and reproducible method (IN-Seq) to fractionate cardiac tissue for proteomic applications [12, 17, 18]. The IN-Seq method sequentially produces three subcellular fractions based on protein solubility at different pH: (1) cytoplasmic-enriched extract (neutral pH), (2) sarcomeric-enriched extract (acidic pH), and (3) membrane protein-enriched pellet (neutral pH). This method was developed to deplete the high abundant sarcomeric sub-proteome, allowing enhanced observation of the cytosolic proteins, while preserving the original endogenous PTM status [17]. In all these sample fractionation methods, each fraction is analyzed separately by nano liquid chromatography– tandem mass spectrometry (nanoLC–MS/MS) for increased total peak capacity while balancing data acquisition time to increase proteome coverage.

While sample fractionation prior to LC-MS is an effective method to improve proteomic depth, the protocol is not easily adapted to high-throughput processing, as it requires extensive manual sample handling and multiple fractions need to be analyzed [19]. Thus, it would be optimal if the fractionation of the peptides can take place in the gas phase [20]. High field asymmetric waveform ion mobility spectrometry (FAIMS) was developed in recent years with the goal of solving fractionation reproducibility [21, 22]. This technology allows for rapid and effective gas-phase separation of peptide ions as they depart the electrospray emitter and prior to their entrance into the mass spectrometer [23]. Combining multiple compensation voltages (CVs) within a single run or between runs improves whole proteome coverage in cell lysates [24, 25]. One example that Pfammatter *et. al*. observed a 30% gain in unique peptide identification using FAIMS compared with non-FAIMS analysis of HEK 293 cells [26]. Different studies using HeLa protein digest such as Johnson KR *et. al*. [27], Greguš M *et. al*. [28], and Cong Y *et. al*. [29], concurrently showed FAIMS had significantly increased in protein and peptide identifications than without FAIMS. In a few recent studies working with various tissue samples, including brain autopsies [30], tumor biopsies [31], or paraffin embedded tissues (lymph node, lung, and prostate) [32], the addition of FAIMS also showed enhanced detection of non-redundant proteoforms, increased proteome sensitivity, or substantially reduced sample handling. As the cardiac proteome consists of large dynamic range, it remains unclear whether FAIMS can detect peptides and the less abundant proteins from the cardiac proteome that is comparable to the previous established fractionation methods.

The total proteomic workflow is comprised of multiple experimental components: sample preparation, data acquisition by LC-MS/MS, and data analysis. Here we present a thorough comparison of the two methods for cardiac muscle tissue (n=3 different hearts): manual IN-Seq fractionation and FAIMS ion separation approach by testing them on the same heart tissue samples split in half for each method preparation. Switching IN-Seq sample fractionation to FAIMS gas-phase separation reduces manual sample handling and shortens MS instrument utilization time to only one third (360 minutes to 120 minutes with FAIMS). The proteome coverage and sensitivity, and peptide intensity between the IN-Seq and FAIMS methods are also compared using data-independent acquisition-MS (DIA-MS) with the same mouse peptide library. We conclude that both methods produce comparable proteomics data, with the FAIMS method proving advantageous for studies requiring higher throughput and efficiency, while IN-Seq provides broader coverage that could be leveraged for in-depth hypothesis generating studies.

## Material and Methods

### Sample Collection

Adult wild-type mice (C57BL/6, n = 3) were sacrificed, and the hearts were immediately harvested, washed in cold PBS buffer. Left ventricles (LV) of each heart was dissected and snap-frozen in liquid nitrogen. LVs were cut in half for “IN-Seq” or “FAIMS” preparations separately (Figure 1, figure created with Biorender.com).

**Fig 1:**
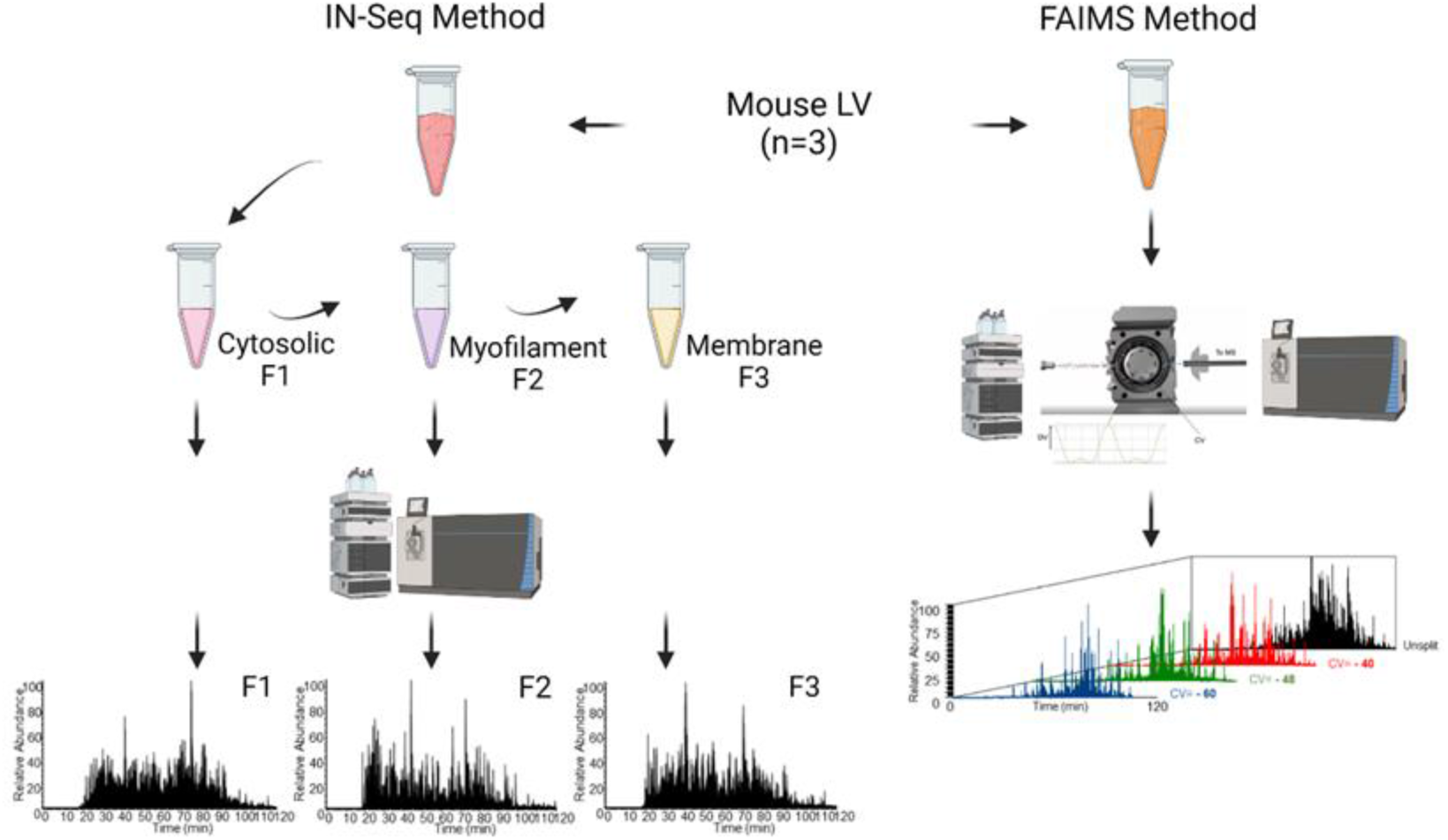
Experimental workflow from the frozen heart tissues to proteomic data analysis. IN-Seq fractionation method (left): Heart tissues (n=3) are homogenized in HEPES extraction buffer, then manually fractionated into cytoplasmic-protein fraction (F1), sarcomeric-protein fraction (F2), and membrane-protein fraction (F3). Total of 3 fractions are processed and prepared separately for mass spectrometry analysis (DIA-MS, Orbitrap Fusion Lumos MS, ThermoFisher). Raw MS data from each fraction are converted to MzML file for DIANN library free analysis. Each tissue sample takes average 1 hour for the manual preparation step and 6 hours of additional MS processing time (3 x120-minute gradient). Field Asymmetric Ion Mobility Spectrometry (FAIMS) Method (right): Tissues from the same hearts (n=3) are homogenized in 8M Urea extraction buffer. Total lysate is directly processed and prepared for mass spectrometry analysis (DIA-MS, Orbitrap Fusion Lumos MS, with FAIMS, ThermoFisher). Raw MS data from the sample is split by compensation voltages (CV-40, CV-48, CV-60) and then converted to MzML files for DIANN library free analysis. MS processing time per tissue sample is about 15 minutes manual preparation and 2 hours of additional MS processing time (120-minute gradient).

### IN-Seq method: sample preparation with sequential fractionation and LC-MS/MS proteomics analysis

Frozen LV halves were minced in ice-cold HEPES extraction buffer (25 mM HEPES-NaOH, 2.5 mM EDTA, pH 7.8) supplemented with Pierce protease inhibitor mini tablets (EDTA free, Thermo Fisher Scientific, Waltham, MA) and Pierce phosphatase inhibitor mini tablets (Thermo Fisher Scientific, Waltham, MA). Samples were fractionated into cytosolic-, sarcomeric-, and membrane protein-enriched fractions (*i*.*e*., F1, F2, F3) by the previously described “IN-Seq” method [12, 33]. Briefly, each tissue sample was homogenized three times in HEPES extraction buffer for 1 minute with a frequency of 30/s. Following centrifugation at 14,000 g for 10 minutes at 4°C. Supernatant was collected and kept at 4°C as F1. The pellet was resuspended in TFA extraction buffer (1% trifluoroacetic acid (TFA), pH 2, 1mM TCEP in HEPES extraction buffer) and homogenized three times for 1minute with at frequency of 30/s. Following centrifugation at 14,000 g for 10 minutes at 4°C, and supernatant was collected and kept at 4°C as F2. Insoluble pellet from F2 extraction step was further solubilized in the SDS extraction buffer (2% SDS, 1mM TCEP in HEPES extraction buffer) and homogenized at frequency of 30/s for 1 minutes, following sonication with a Q800R3 Sonicator (QSonica, Newtown, CT) for 5 minutes (amplitude of 70%, with 10s on, 10s off) and then centrifugation in 14,000g for 10 minutes at 4°C, collecting the supernatant as F3. Protein concentration of all fractions of each sample were determined by BCA (Pierce, Waltham, MA). 100 μg of each fraction (total protein quantification) were reduced using 1 mM TCEP. Clean-up of each fraction was performed using S-trap mini spin columns (ProtiFi, Long Island, NY) for complete destruction of undesired enzymatic activity (such as protease or phosphatase activities) and maximization of digestion efficiency. Each fraction was digested for 15–18h at 37°C using ultra-grade Trypsin (Promega) at a 1:100 enzyme: protein ratio. All fractions from each tissue samples were speed-vacuum dried and stored at -80°C until MS analysis.

For DIA-MS analysis with the IN-Seq fractionated samples, we used the Orbitrap Lumos Fusion mass spectrometer (Thermo Scientific) equipped with an EasySpray ion source and connected to Ultimate 3000 nano LC system (Thermo Scientific) as previously described [33]. Peptides were loaded onto a PepMap RSLC C18 column (2 μm, 100 Å, 150 μm i.d. x 15 cm, Thermo) using a flow rate of 1.4 μL/min for 7 minutes at 1% B (mobile phase A was 0.1% formic acid in water and mobile phase B was 0.1 % formic acid in acetonitrile) after which point they were separated with a linear gradient of 5-20%B for 45 minutes, 20-35%B for 15 minutes, 35-85%B for 3 minutes, holding at 85%B for 5 minutes and re-equilibrating back to 1% B over the course of 5 minutes. For the longer 120-minute gradient method, peptides were separated with a linear gradient of 5-20%B for 90 minutes, 20-35% for 30 minutes, 35-85%B for 3 minutes, holding at 85%B for 5 minutes and re-equilibrating back to 1% B over the course of 5 minutes. Each fraction was followed by a blank injection to both clean the column and re-equilibrate at 1%B. The nano-source capillary temperature was set to 300 °C and the spray voltage was set to 1.8 kV. Indexed retention time standards (iRTs, Biognosys) were added to each fraction before acquisition [34].

### FAIMS method: tissue sample preparation and LC-FAIMS-MS/MS proteomics analysis

The other frozen halves of LV samples were subjected to polytron homogenization and denaturation in UREA total protein extraction buffer (8M Urea, 1M Ammonium Bicarbonate, 5% SDS) at 4°C. Tissue homogenates were ultrasonicated with a Q800R3 Sonicator (QSonica, Newtown, CT) for 5 minutes (amplitude of 70%, with 10s on, 10s off) and then centrifuged in 20,000g for 10 minutes at 4°C. Supernatant of each sample was transferred and collected as the total protein extracts. Protein concentration of each tissue sample was determined by BCA (Pierce, Waltham, MA). S-trap mini spin columns (ProtiFi, Long Island, NY) was used to clean up 100 μg of each tissue sample, following ultra-grade Trypsin (Promega) digestion at a 1:100 enzyme: protein ratio for 15-18h at 37°C. All tissue samples were dried using speed vacuum and stored at -80°C until MS analysis.

DIA-MS was carried out using established protocols on the Orbitrap Lumos Fusion MS (the same instrument as In-Seq) with a FAIMS front end (Thermo) coupled to a stainless-steel emitter (EvoSep) and EasySpray source (Thermo) adapter [35]. The trapping column (75 µm ID x length 3 µm Luna particles, Phenomenex) was run with 0.1% aqueous formic acid for 7 minutes at 5 µL/min. Samples were diverted onto a 200 cm micro pillar array column (µPAC, PharmaFluidics) and peptides separated using a gradient of mobile phases A and B composed of 0.1% aqueous formic acid and 0.1% formic acid in acetonitrile, respectively. The gradient consisted of 4% B for the first 5 minutes with a step increased to 8% at 5.2 minutes followed by a 90-minute linear gradient up to 30% B. During this time, the flow rate, which started at 1200 nL/min, was linearly decreased to 1000 nL/min. Subsequently, the %B was linearly increased from 30% to 50% over 30 minutes while being held at a constant flow rate of 1000 nL/min for a total gradient run time of 120 minutes. Following each analysis, a 20-minute equilibration was performed, during which the trap was back-flushed at 5 µL/min while the analytical column was washed with 95% B and equilibrated to 2% B at 1200nL/min. Both separation and equilibration were carried out at 55°C.

The FAIMS module was used to separate the electro sprayed peptides by collisional cross section into three populations by cycling the compensation voltage (CV) between -40, -48, and - 60. At each compensation voltage a precursor scan spanning 400-1000 m/z was acquired at 60,000 (at m/z=200) resolution with AGC target set to 1000000 and 50 ms maximum injection time followed by data independent acquisition (DIA) spanning the same range using 20 m/z wide windows (30 windows at each FAIMS CV). Each DIA scan was acquired at 15,000 resolutions (at m/z=200) with AGC set to 150000 and 35 ms maximum injection time.

### Proteomics data processing and analysis

Raw MS/MS data files were converted to mzML format using MSconvert version v.3.0.21304 from ProteoWizard for generating peak lists [36]. The MS files were analyzed in DIA-NN 1.8.1 using the spectral library-free search with Prosit against the Mouse Uniprot database [37]. In the library-free search, neural network algorithms are used to accurately predict hypothetical spectra and retention times for each potential tryptic peptide in the database. Only proteotypic peptides that consist of amino acid sequences unique to one proteins are used for protein identification [38]. The search was conducted with the second pass and match between runs (MBR) options enabled. The quantitation at the protein levels was summed between the different CV fractions. MS protein quantification for the IN-Seq method samples was carried out by averaging the raw peptide intensity among technical replicates and summed among three sub-cellular fractions. Statistical analysis and software data was filtered by interquartile range and normalized to the median. One-way ANOVA with post-hoc Fisher’s LSD test was used to compare datasets with more than two groups. An unpaired t-test was used to analyze the rest of the datasets. A p value of ≤0.05 was considered significant. MetaboAnalyst 5.0 was used for statistical analysis and data visualization [39]. Cellular component visualization was performed using Protein Interaction Network Extractor (PINE) [40].

## Results and Discussion

### FAIMS method shortens sample preparation time and increases instrument efficiency for proteomics analysis

The time of sample preparation step(s) between FAIMS method and the IN-Seq method. FAIMS method requires ∼15 minutes per sample (for 5 minutes of tissue homogenization, 10 minutes of tissue solubilization and protein denaturation) before proteolytic digestion. In contrast, IN-Seq method requires additional steps for the manual fractionation of each sample into the three subcellular fractions which involves addition of multiple buffers, homogenization, and centrifugation steps. This takes ∼1 hour per heart tissue sample (for tissue homogenization, tissue solubilization, protein denaturation and fractionation) before proteolytic digestion. Moreover, each fraction gets injected separately into the LC-MS/MS with the same 120-minute gradient for proteomics analysis, which makes instrument processing time 360 minutes per heart sample (6 hrs), while total LC-FAIMS-MS/MS time using the same gradient, takes only 120 minutes (2 hrs).

FAIMS allows for a larger number of samples to be processed per day and reduces potential batch effects when processing large number of samples [41]. It shortens instrument time, reduces the time between data acquisition and analysis. Finally, FAIMS approach lowers the use of reagents during sample preparation compared to the IN-Seq method.

### FAIMS method covers the majority of proteome identified by IN-Seq method

We next investigated whether FAIMS method produces comparable proteome coverage compared to tissue analyzed with the IN-Seq method. Total of 2211 proteins were identified using the FAIMS method while 2771 proteins from all three fractions using the IN-Seq method. There was an overlap of 2172 proteins, which account for 98% of the total proteome identified by FAIMS and 78% of the total proteome by IN-Seq. 599 proteins (22% of total proteins) were identified uniquely by the IN-Seq method, while 39 proteins (2% of total proteins) were uniquely identified by FAIMS (Figure 2A). For protein intensity distributions, IN-Seq detected more proteins in the lower intensity range; however, with the common 2172 proteins identified by both methods, FAIMS exhibited relatively higher protein intensity (Figure 2B, C). For the proteins uniquely identified by each method (599 from IN-Seq method vs. 39 from FAIMS method), based on cellular component analysis using Protein Interaction Network Extractor (PINE), IN-Seq method maintained high sensitivity for detecting cytoplasmic proteins, especially intracellular membrane-bounded organelle proteins, compared to FAIMS method.

**Fig 2.**
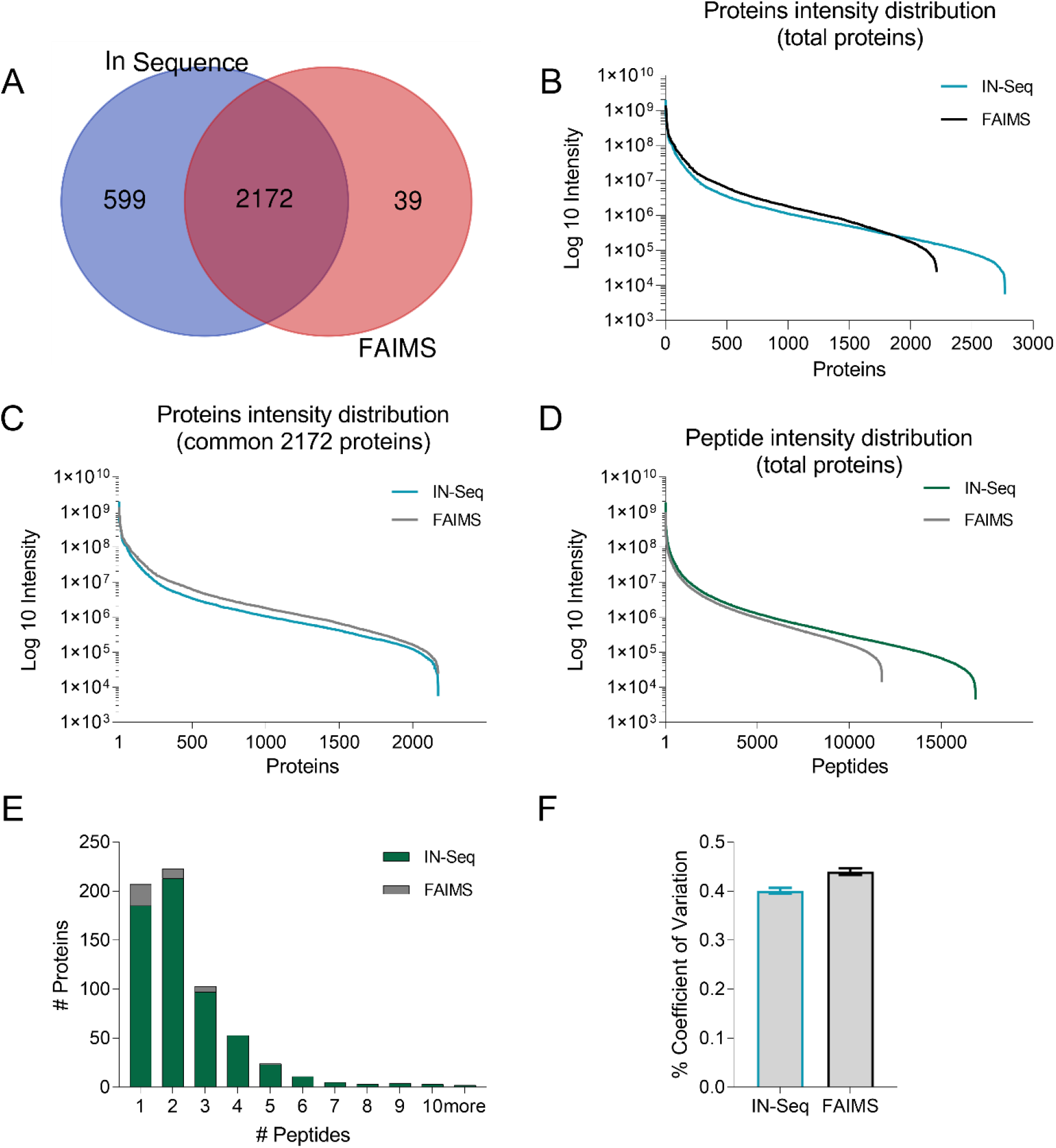
Proteomic data comparing the IN-Seq fractionation method and FAIMS method. A. Venn diagram of proteins identified by each method and the total protein overlap. B. Peak intensity distribution of the common 2172 proteins identified by both methods. C. Peak intensity distribution of the 2771 proteins identified by IN-Seq fractionation method and the 2211 proteins identified by the FAIMS method. D. Peak intensities of all peptides identified by the In Sequence method and the FAIMS method, peptide intensity from the most to least intense. E. Numbers of peptides for the uniquely identified proteins by the IN-Seq method (599 proteins), and uniquely identified by the FAIMS method (39 proteins). F. %CV between IN-Seq and FAIMS methods.

To understand if there was a bias in the precursor peptides, we further investigated the peptide level data after the DIA-NN proteomics analysis for sensitivity and reproducibly. It is important to realize that only proteolytic peptides, peptides comprised of amino acid sequence that is unique to a protein are included. IN-Seq method has slightly higher sensitivity in detecting peptides comparing to the FAIMS methods (Figure 2D). The FAIMS unique proteins mostly had one and two proteotypic peptide to one protein ratio and greater numbers than In-Seq. While the IN-Seq unique proteins had more two proteotypic peptides to one protein ratio which is helpful when screening for PTMs (Figure 2E). This means that with common criteria for protein quantification of 2 prototypic peptides that FAIMs is like IN-Seq, especially as the two methods have comparable percentage of coefficient of variance (%CV, Figure 2F) with majority of peptides having CV% under 40% based on 3 biological replicates.

## Conclusion

In the present study, we examined whether LC-FAIMS-MS/MS in ion gas-phase separation can be adapted as the alternative method for heart tissue proteomics study compared to the commonly used IN-Seq fractionation LC-MS/MS method. Using the same heart samples (n=3) between the two methods, FAIMS detected over 79% of IN-Seq detected proteins, with one third reagent use in tissue preparation and much shortened MS instrument time. IN-Seq method remained to be ideal for studies focused on changes in cytosolic proteome of cardiac tissues, as well as studies involving smaller sample numbers. Both methods produced comparable results of cardiac proteome profiles, but the FAIMS method presented here provided much improved throughput and shortened MS processing time compared to the IN-Seq method.

**Table 1:** Comparison between FAIMS and IN-Seq Methods. Processing time and proteome coverage breakdown for the IN-Seq method and the FAIMS method.

| Method | Tissue Processing Time | Instrument Time | Identified Proteins | Unique Identified Proteins |
| --- | --- | --- | --- | --- |
| FAIMS | 15 min | 120 min | 2211 | 39 |
| IN-Seq | 60 min | 360 min | 2771 | 599 |

## Reference

1. Smith, R.D., Y. Shen, and K. Tang, Ultrasensitive and Quantitative Analyses from Combined Separations−Mass Spectrometry for the Characterization of Proteomes. Accounts of Chemical Research, 2004. 37(4): p. 269–278.

2. Arrell, D.K., I. Neverova, and J.E.V. Eyk, Cardiovascular Proteomics. 2001. 88(8): p. 763–773.

3. McGregor, E. and M.J. Dunn, Proteomics of the Heart. 2006. 98(3): p. 309–321.

4. de Castro Brás, L.E., et al., Texas 3-step decellularization protocol: looking at the cardiac extracellular matrix. J Proteomics, 2013. 86: p. 43–52.

5. Chugh, S., C. Suen, and A. Gramolini, Proteomics and mass spectrometry: what have we learned about the heart? Curr Cardiol Rev, 2010. 6(2): p. 124–33.

6. Aballo, T.J., et al., Ultrafast and Reproducible Proteomics from Small Amounts of Heart Tissue Enabled by Azo and timsTOF Pro. J Proteome Res, 2021. 20(8): p. 4203–4211.

7. Gramolini, A.O., et al., Analyzing the cardiac muscle proteome by liquid chromatography-mass spectrometry-based expression proteomics. Methods Mol Biol, 2007. 357: p. 15–31.

8. Cammarato, A., et al., A mighty small heart: the cardiac proteome of adult Drosophila melanogaster. PLoS One, 2011. 6(4): p. e18497.

9. Lau, E., et al., A large dataset of protein dynamics in the mammalian heart proteome. Scientific Data, 2016. 3(1): p. 160015.

10. White, M.Y., et al., Parallel Proteomics to Improve Coverage and Confidence in the Partially Annotated <em>Oryctolagus cuniculus</em> Mitochondrial Proteome *<sup> </sup>. Molecular & Cellular Proteomics, 2011. 10(2): p. S1–S15.

11. Doll, S., et al., Region and cell-type resolved quantitative proteomic map of the human heart. Nature Communications, 2017. 8(1): p. 1469.

12. Kane, L.A., I. Neverova, and J.E. Van Eyk, Subfractionation of Heart Tissue, in Cardiovascular Proteomics: Methods and Protocols, F. Vivanco, Editor. 2007, Humana Press: Totowa, NJ. p. 87–90.

13. Lizotte, E., et al., Isolation and characterization of subcellular protein fractions from mouse heart. Analytical Biochemistry, 2005. 345(1): p. 47–54.

14. Jones, L.R., et al., Separation of vesicles of cardiac sarcolemma from vesicles of cardiac sarcoplasmic reticulum. Comparative biochemical analysis of component activities. Journal of Biological Chemistry, 1979. 254(2): p. 530–539.

15. Boivin, B. and B.G. Allen, Regulation of membrane-bound PKC in adult cardiac ventricular myocytes. Cellular Signalling, 2003. 15(2): p. 217–224.

16. Warren, C.M., et al., Sub-proteomic fractionation, iTRAQ, and OFFGEL-LC-MS/MS approaches to cardiac proteomics. J Proteomics, 2010. 73(8): p. 1551–61.

17. Agnetti, G., C. Husberg, and J.E.V. Eyk, Divide and Conquer. 2011. 108(4): p. 512–526.

18. Neverova, I., Role of chromatographic techniques in proteomic analysis. Journal of chromatography. B, Analytical technologies in the biomedical and life sciences. 815(1-2): p. 51–63.

19. Swearingen, K.E. and R.L. Moritz, High-field asymmetric waveform ion mobility spectrometry for mass spectrometry-based proteomics. Expert Rev Proteomics, 2012. 9(5): p. 505–17.

20. Cooper, H.J., To What Extent is FAIMS Beneficial in the Analysis of Proteins? J Am Soc Mass Spectrom, 2016. 27(4): p. 566–77.

21. Krylov, E.V. and E.G. Nazarov, Electric field dependence of the ion mobility. International Journal of Mass Spectrometry, 2009. 285(3): p. 149–156.

22. Schneider, B.B., et al., Planar differential mobility spectrometer as a pre-filter for atmospheric pressure ionization mass spectrometry. Int J Mass Spectrom, 2010. 298(1-3): p. 45–54.

23. Hebert, A.S., et al., Comprehensive Single-Shot Proteomics with FAIMS on a Hybrid Orbitrap Mass Spectrometer. Analytical Chemistry, 2018. 90(15): p. 9529–9537.

24. Schweppe, D.K., et al., Characterization and Optimization of Multiplexed Quantitative Analyses Using High-Field Asymmetric-Waveform Ion Mobility Mass Spectrometry. Analytical Chemistry, 2019. 91(6): p. 4010–4016.

25. Schweppe, D.K., et al., Optimized Workflow for Multiplexed Phosphorylation Analysis of TMT-Labeled Peptides Using High-Field Asymmetric Waveform Ion Mobility Spectrometry. Journal of Proteome Research, 2020. 19(1): p. 554–560.

26. Pfammatter, S., et al., A Novel Differential Ion Mobility Device Expands the Depth of Proteome Coverage and the Sensitivity of Multiplex Proteomic Measurements. Mol Cell Proteomics, 2018. 17(10): p. 2051–2067.

27. Johnson, K.R., M. Greguš, and A.R. Ivanov, Coupling High-Field Asymmetric Ion Mobility Spectrometry with Capillary Electrophoresis-Electrospray Ionization-Tandem Mass Spectrometry Improves Protein Identifications in Bottom-Up Proteomic Analysis of Low Nanogram Samples. J Proteome Res, 2022. 21(10): p. 2453–2461.

28. Greguš, M., et al., Improved Sensitivity of Ultralow Flow LC–MS-Based Proteomic Profiling of Limited Samples Using Monolithic Capillary Columns and FAIMS Technology. Analytical Chemistry, 2020. 92(21): p. 14702–14712.

29. Cong, Y., et al., Ultrasensitive single-cell proteomics workflow identifies >1000 protein groups per mammalian cell. Chemical Science, 2021. 12(3): p. 1001–1006.

30. Fulcher, J.M., et al., Enhancing Top-Down Proteomics of Brain Tissue with FAIMS. Journal of Proteome Research, 2021. 20(5): p. 2780–2795.

31. Sweet, S., et al., The addition of FAIMS increases targeted proteomics sensitivity from FFPE tumor biopsies. Scientific Reports, 2022. 12(1): p. 13876.

32. Eckert, S., et al., Evaluation of Disposable Trap Column nanoLC-FAIMS-MS/MS for the Proteomic Analysis of FFPE Tissue. J Proteome Res, 2021. 20(12): p. 5402–5411.

33. Soetkamp, D., et al., Myofilament Phosphorylation in Stem Cell Treated Diastolic Heart Failure. 2021. 129(12): p. 1125–1140.

34. Robinson, A.E., et al., Lysine and Arginine Protein Post-translational Modifications by Enhanced DIA Libraries: Quantification in Murine Liver Disease. Journal of Proteome Research, 2020. 19(10): p. 4163–4178.

35. Plummer, J.T., et al., US Severe Acute Respiratory Syndrome Coronavirus 2 (SARS-CoV-2) Epsilon Variant: Highly Transmissible but With an Adjusted Muted Host T-Cell Response. Clinical Infectious Diseases, 2022.

36. Chambers, M.C., et al., A cross-platform toolkit for mass spectrometry and proteomics. Nature Biotechnology, 2012. 30(10): p. 918–920.

37. Demichev, V., et al., DIA-NN: neural networks and interference correction enable deep proteome coverage in high throughput. Nature Methods, 2020. 17(1): p. 41–44.

38. Craig, R., J.P. Cortens, and R.C. Beavis, The use of proteotypic peptide libraries for protein identification. Rapid Commun Mass Spectrom, 2005. 19(13): p. 1844–50.

39. Pang, Z., et al., Using MetaboAnalyst 5.0 for LC–HRMS spectra processing, multi-omics integration and covariate adjustment of global metabolomics data. Nature Protocols, 2022. 17(8): p. 1735–1761.

40. Sundararaman, N., et al., PINE: An Automation Tool to Extract and Visualize Protein-Centric Functional Networks. J Am Soc Mass Spectrom, 2020. 31(7): p. 1410–1421.

41. Bekker-Jensen, D.B., et al., A Compact Quadrupole-Orbitrap Mass Spectrometer with FAIMS Interface Improves Proteome Coverage in Short LC Gradients*. Molecular & Cellular Proteomics, 2020. 19(4): p. 716–729.

