## Supplementary Material 5 for "High Field Asymmetric Waveform Ion Mobility Spectrometry-Mass Spectrometry to Enhance Cardiac Muscle Proteome Coverage"

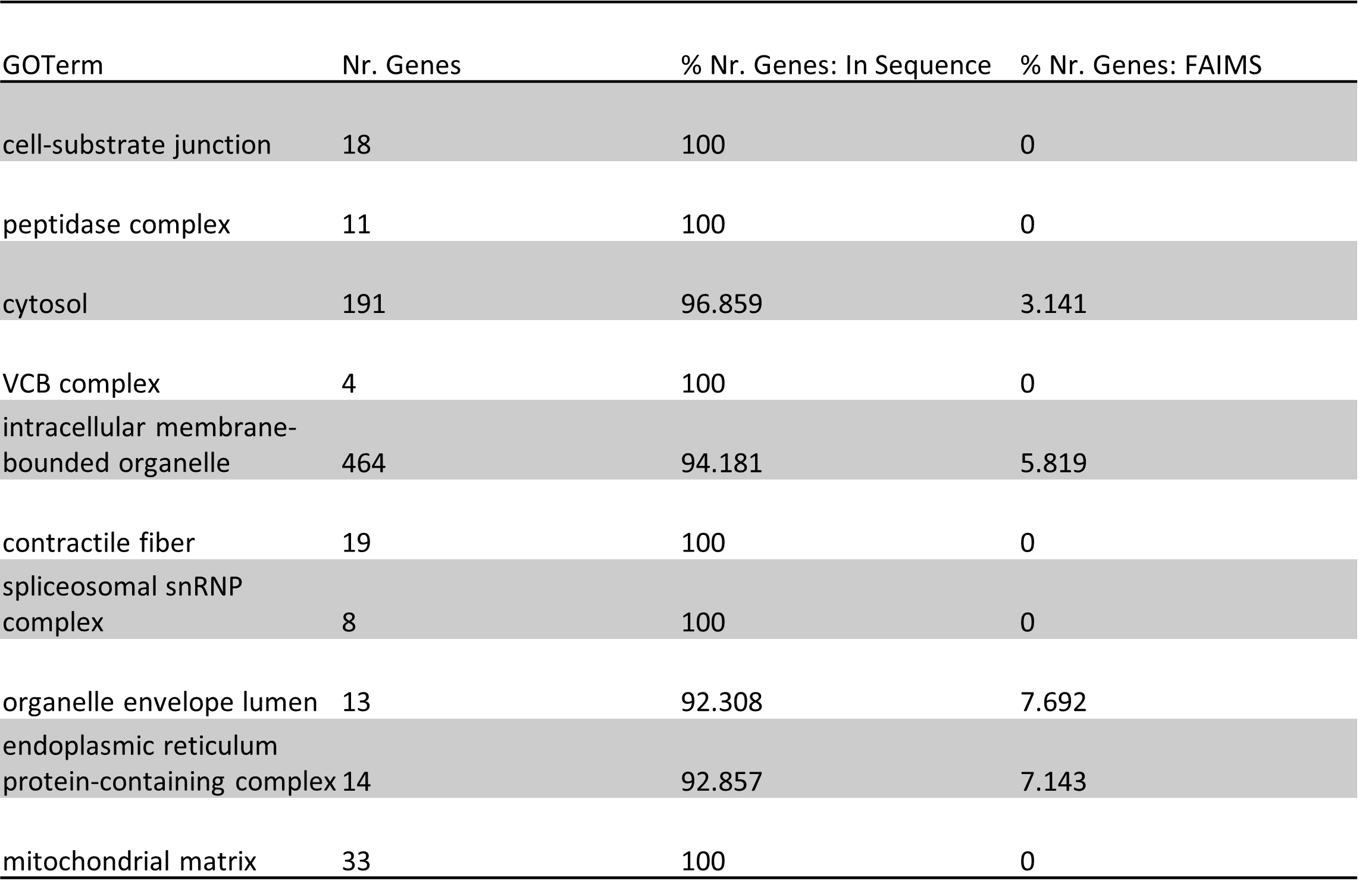

Supplementary table: GO Terms – Cellular Component percentage analysis comparing between the 39 proteins that uniquely identified by FAIMS method and the 599 proteins that uniquely identified by the IN-Seq method.
